# Striatonigral distribution of a fluorescent reporter following intracerebral delivery of genome editors

**DOI:** 10.1101/2023.06.21.542185

**Authors:** Samuel S. Neuman, Jeanette M. Metzger, Viktoriya Bondarenko, Yuyuan Wang, Jesi Felton, Jon E. Levine, Krishanu Saha, Shaoqin Gong, Marina E. Emborg

**Affiliations:** Wisconsin National Primate Research Center, University of Wisconsin-Madison, Madison, WI, USA; Department of Ophthalmology and Visual Sciences, University of Wisconsin–Madison, Madison, WI, USA; Department of Biomedical Engineering, University of Wisconsin–Madison, Madison, WI, USA; Wisconsin Institute for Discovery, University of Wisconsin–Madison, Madison, WI, 53715, USA; Department of Neuroscience, University of Wisconsin–Madison, Madison, WI, USA; Department of Medical Physics, University of Wisconsin-Madison, Madison, WI, USA

**Keywords:** Nanomedicine, CRISPR, Viral Vectors, Parkinson’s Disease, Neural Networks, Substantia Nigra, Gene Editing

## Abstract

**Introduction:** Targeted gene editing is proposed as a therapeutic approach for numerous disorders, including neurological diseases. As the brain is organized into neural networks, it is critical to understand how anatomically connected structures are affected by genome editing. For example, neurons in the substantia nigra pars compacta (SNpc) project to the striatum, and the striatum contains neurons that project to the substantia nigra pars reticulata (SNpr).

**Methods:** Here, we report the effect of injecting genome editors into the striatum of Ai14 reporter mice, which have a LoxP-flanked stop cassette that prevents expression of the red fluorescent protein tdTomato. Two weeks following intracerebral delivery of either synthetic nanocapsules (NCs) containing CRISPR ribonucleoprotein targeting the tdTomato stop cassette or adeno-associated virus (AAV) vectors expressing Cre recombinase, the brains were collected, and the presence of tdTomato was assessed in both the striatum and SN.

**Results:** TdTomato expression was observed at the injection site in both the NC- and AAV-treated groups and typically colocalized with the neuronal marker NeuN. In the SN, tdTomato-positive fibers were present in the pars reticulata, and SNpr area expressing tdTomato correlated with the size of the striatal genome edited area.

**Conclusion:** These results demonstrate *in vivo* anterograde axonal transport of reporter gene protein products to the SNpr following neuronal genome editing in the striatum.

## 1 Introduction

*In vivo* somatic gene editing is currently being researched as a therapy for numerous diseases, including neurological disorders (Heidenreich and Zhang, 2016; Saha et al., 2021; Sharma et al., 2021). Delivery of genome editors, such as CRISPR Cas9, directly into the brain has the potential to allow for targeted correction of disease-associated mutations in specific neuronal populations. However, no neuronal population exists in isolation. The network-like composition of the brain makes it critical to understand how genome editing in a given region will impact anatomically connected structures.

One example of a neural network in the brain is the nigrostriatal system (Dahlstroem and Fuxe, 1964; Björklund and Dunnett, 2007; Poulin et al., 2018; Harris et al., 2020). The substantia nigra (SN) is located in the ventral midbrain and consists of the substantia nigra pars compacta (SNpc) and pars reticulata (SNpr). The dopaminergic neurons in the SNpc provide innervation to the GABAergic medium spiny neurons of the dorsal striatum. The striatum, located at the core of the forebrain, is a complex, heterogenous structure with multiple subregions. Projections from the striatum reach many brain areas, and a key striatal output is the SNpr. Thus, a two-way connection exists between the striatum and the SN.

Neurons in the SNpc are progressively lost with aging, and excessive, accelerated loss of dopaminergic nigral neurons leads to the development of Parkinson’s disease (PD) (Fearnley and Lees, 1991; Lee et al., 1995; Emborg et al., 1998; Baldereschi et al., 2000; Chu and Kordower, 2007). Surgical delivery of prospective neuroprotective strategies, such as gene editors, to the SN is difficult due to its small size, location deep within the brain, and characteristically vulnerable neuronal population in PD (Albert et al., 2017). Targeting of the striatum for genome editor delivery could impact protein expression in the SN through either anterograde transport along axons of striatal neurons projecting to the SNpr or retrograde transport of editors taken up by axonal terminals of SNpc neurons.

Recently, our research group achieved successful gene editing of striatal neurons using a novel CRISPR-Cas9 ribonucleoprotein (RNP) nanocapsule (NC). This NC features glutathione-responsive crosslinkers and imidazole-containing monomers capable of endosomal escape, creating a delivery vehicle that is stable, yet biodegradable, that can release RNP upon entering a cell (Chen et al., 2019). NC delivery of CRISPR RNP produced robust and efficient editing of multiple striatal neuron types, including GABAergic neurons (Metzger et al., 2023). However, the impact of this NC-induced striatal genome editing on neural networks of the brain has not been assessed.

In this study, we investigated delivery of both the novel CRISPR Cas9 RNP NCs and adeno-associated virus vector serotype 9 (AAV9) as neural network strategies for neuronal gene editing. These genome editors were stereotactically injected in the striatum of Ai14 reporter mice. Expression of the reporter transgene tdTomato was then assessed at the site of injection in the striatum as well as the SNpc and SNpr.

## 2 Materials and Methods

### 2.1 Intrastriatal delivery of genomic editing material

Cas9-sgRNA ribonucleoproteins (RNPs) were prepared according to previously published protocols (Metzger et al., 2023). Nuclear localization signal (NLS)-tagged Streptococcus pyogenes Cas9 nuclease (sNLS-SpCas9-sNLS) was obtained from Aldevron and combined and complexed at a 1:1 molar ratio with sgRNA (Integrated DNA Technologies) (Wang et al., 2021). The sgRNAs used in this experiment include the Ai14 guide RNA (sgRNA) (protospacer 5′ - AAGTAAAACCTCTACAAATG-3′) and a non-targeting sgRNA (Alt-R CRISPR-Cas9 Negative Control crRNA #1, Integrated DNA Technologies, Inc., USA). NCs were prepared as previously reported (Metzger et al., 2023). AAV9 vectors containing CMV-driven Cre plasmids were obtained from Addgene (pENN.AAV.CMVs.Pl.Cre.rBG #105537). Vectors were stored in a solution containing PBS and 0.001% Pluronic F-68.

Intracerebral injections of NCs or AAV vectors followed previously published protocols (Metzger et al., 2023). All animal treatments and procedures were approved by the University of Wisconsin– Madison Animal Care and Use Committee. Prior to surgery, mice were anaesthetized by intraperitoneal injection of a ketamine (120 mg/kg), xylazine (10 mg/kg), and acepromazine (2 mg/kg) cocktail. Ai14 mice (The Jackson Laboratory, STOCK# 7914) underwent stereotactic brain injections at coordinates of AP +0.74 mm, ML ±1.82 mm, DV -3.37 mm in the striatum, enabling assessment of potential retrograde and anterograde transport to the SN (Figure 1A,B; Figure created with BioRender.com). Injections were performed via a Stoelting stereotaxic frame equipped with a Stoelting Quintessential Stereotax Injector (QSI). Intracerebral delivery occurred at a rate of 0.2 μl/minute using a 10 μL Hamilton syringe and 32-gauge 1 inch Hamilton small hub RN needle. The NC delivery included 1.5 μL injection of NC carrying 20 μM of RNP guide targeting Ai14 (n=9) or a non-targeting guide (n=9) in corresponding NC storage buffer as previously described (Metzger et al., 2023). The AAV delivery included 1 uL of ≥ 1×10^13^ vg/mL titer (n=4) in corresponding, as-prepared storage buffer. Following completion of the injection, the needle remained in place for up to 5 minutes. The surgical field was then irrigated with sterile saline, and the skin layers were closed using surgical glue.

**Figure 1.**
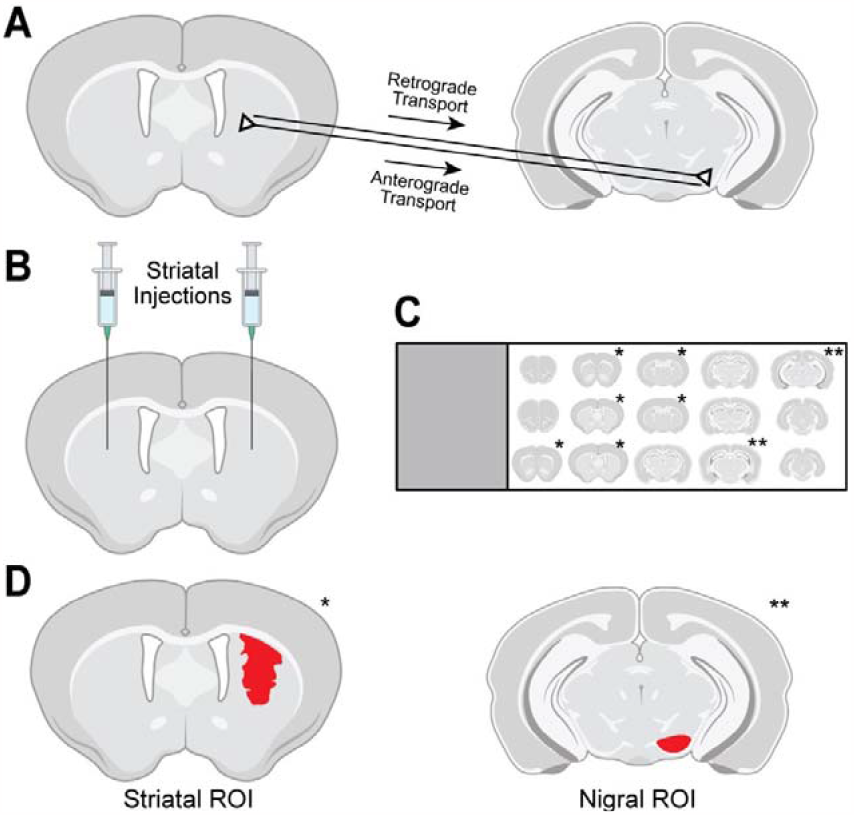
Methods used for evaluation of tdTomato expression in the striatum and substantia nigra (SN) of Ai14 mice. (A) Following intrastriatal injection of genome editors, tdTomato in the SN pars compacta (SNpc) results from retrograde transport of editors from axon terminals to SNpc soma and subsequent gene editing. Expression in the SN pars reticulata (SNpr) results from anterograde transport of tdTomato protein from successfully edited striatal neurons along their axons. (B) Intrastriatal injections in Ai14 mice delivered adeno-associated virus (AAV) vectors encoding Cre recombinase or CRISPR nanocapsules (NCs). (C) After necropsy, serial tissue sections 240 μm apart spanning the rostrocaudal spread of the brain were immunostained and mounted on slides, and sections containing either striatum (*) or SN (**) were imaged using confocal microscopy. (D) For analysis, regions of interest (ROIs; red) were drawn in the striatum around all tdTomato-expressing cells and in the SN around the anatomically defined pars reticulata or pars compacta.

### 2.2 Necropsy and Tissue Processing

Fourteen days after surgery, mice were deeply anesthetized with a combination of ketamine (120 mg/kg) and xylazine (10 mg/kg) and were transcardially perfused with heparinized saline. Brains were retrieved, post-fixed for 24 hours in 4% PFA, and cryoprotected in graded sucrose. The brains were cut frozen in 40 μm coronal sections on a sliding microtome and stored at -20°C in cryoprotectant solution (1000 ml 1X PBS (pH 7.4), 600 g sucrose, 600 ml ethylene glycol) until immunostaining. The anatomical left hemisphere of each coronal slice was identified by marking two small holes in the cerebral cortex.

### 2.3 Immunohistochemistry

All immunolabeling was performed using one sixth serial brain sections spaced 240 μm apart as previously described (Metzger et al., 2023) (Figure 1C). The sections were incubated with primary antibodies (Mouse anti-NeuN 1:1000, Abcam ab104224, lot GR3386324-1; Rabbit anti-RFP 647 nm 1:5000, Rockland 600-401-379, lot 42872; Sheep anti-TH 1:1000, Invitrogen PA1-4679, lot WH3357994) for 24 hours at room temperature, washed 3 times in dilution media, and then incubated for 2 hours at room temperature with secondary antibodies in PBS (Donkey anti-Mouse 647 nm 1:1000, Abcam ab15107, lot GR3234832-4; Donkey anti-Rabbit 594nm 1:1000, Invitrogen A21207, lot 1668652; Donkey anti-Sheep 488 nm 1:1000, Jackson ImmunoResearch 713-545-003, lot 149534). Tissue was counterstained with DAPI, mounted onto slides, allowed to dry, and cover slipped with Fluor Gel. Negative controls were performed through omission of primary antibodies.

### 2.4 Image Acquisition and Analysis

Image acquisition was performed on all available striatal and nigral tissue sections in one sixth serial brain sections spaced 240 μm apart (Figure 1C) per animal using a Nikon A1R confocal microscope running NIS Elements version 5.20.02 with 405, 488, 561, and 640 wavelength lasers using as previously described (Metzger et al., 2023). Detectors for the 488 and 561 lasers are high sensitivity GaAsP PMTs, while the 405 and 640 lasers use HS PMTs. Confocal laser power and detector sensitivity settings were the same for animals within the NC treatment groups and within the AAV9-Cre treatment group. Due to noticeably higher red channel signal intensity in the striata of AAV9-Cre-injected animals, lower red channel laser power was used when acquiring images from this treatment group compared to NC-Ai14 or NC-Scrambled animals. The 4x objective (Plan Apo, N.A. = 0.2, Nikon) was used to acquire images of whole coronal brain slices. The 20x objective (NA0.75 GLYC WD = 0.66 mm CORR) was used to obtain images of the striatal injection site and substantia nigra via acquisition of three focal planes covering a total of 10 μm, which were processed as maximum intensity projections.

In the striatum, the total size of the tdTomato+ edited area was quantified by drawing regions of interest (ROIs) in 20x images around areas of tdTomato signal in which a group of at least 5 cells were tdTomato+ and within 135 μm of each other (Figure 1D). ROIs were drawn in NIS Elements version 5.30.02 using the Draw Polygonal ROI function in the red (tdTomato) channel, and the size (area in mm^2^) was exported for each ROI. ROIs were drawn below the corpus callosum (visualized with NeuN and DAPI staining). The total striatal genome edited area for each brain hemisphere was then calculated by summing the striatal ROI areas across tissue sections analyzed in that hemisphere.

In the SNpr, the size of the tdTomato+ area was evaluated by drawing ROIs around the complete SNpr according to the Allen Mouse Brain Atlas (Lein et al., 2007) (Figure 1D) and then using ImageJ (version 1.54d) to quantify the area of each ROI with tdTomato. First, using 20x images, the 594 nm (tdTomato) channel was isolated, converted to an 8 bit image, and inverted. Next, the SNpr ROI was drawn, and the ImageJ threshold function was applied to detect tdTomato. To select a threshold to exclude any background in the 594 nm channel, a square ROI was drawn outside of the SNpr, the pixel intensity evaluated in the square ROI, and a pixel intensity threshold cutoff was used just above the background pixel intensity. The size (area in mm^2^) of the area above threshold was exported for each ROI. Due to variations in the number of sections with visible SN across animals, the two SNpr sections in each hemisphere with the largest SNpr tdTomato+ area were summed for analysis.

In the SNpc, ROIs were drawn in NIS Elements with reference to the Allen Mouse Brain Atlas (Lein et al., 2007) and 4x images. The total number of genome edited neurons present in each SNpc ROI was counted as the number of TdTomato+ and NeuN+ cells.

### 2.5 Statistics

All statistical analyses were performed in MATLAB R2022a. Data values in the text are presented as mean ± standard error of the mean. The Pearson’s correlation coefficient was calculated to test the relationship between tdTomato+ area in the striatum and the %AAT in the SNpr. T tests were used for any comparison between treatment groups. All p values reported are two tailed, and a p value < 0.05 is considered statistically significant.

## 3 Results

### 3.1 Successful genome editing was observed in the injected striatum

Genome editing was first confirmed at the striatal injection site in Ai14 mice treated with CRISPR RNP NCs or AAV9-Cre. These mice harbor the Ai14 Cre reporter allele, which contains a Lox-P flanked stop cassette consisting of three SV40 polyA transcriptional terminator sequences upstream of the CAG promoter-driven red fluorescent protein tdTomato. Successful gene editing produces tdTomato expression due to either NC-delivery of CRISPR RNP with guide RNA targeting sequences within the stop cassette (NC-Ai14) leading to removal of at least two SV40 polyA blocks or AAV9-Cre transduction and Cre-Lox recombination.

In both NC-Ai14 and AAV9-Cre treatment groups, abundant tdTomato-immunoreactivity (-ir) was visible in the striatum in coronal brain sections of tissue collected two weeks after stereotactic injection (Figure 2). No tdTomato-ir was observed in animals injected with NCs containing CRISPR RNP with non-targeting guide RNA (NC-Scrambled). The presence of tdTomato was assessed in serial brain sections spaced 240 μm apart to evaluate the total area of gene editing throughout the striatum. In each animal, the injection target site could be identified in the brain section with the largest area of tdTomato-ir, and areas of editing decreased in size in sections rostral and caudal to the target (Figure 2A, B; filled arrowheads). Occasionally, tdTomato-ir was also observed along the needle tract in the cortex superior to the target site (Figure 2A, B; open arrowheads). The total striatal edited area was similar between NC-Ai14 and AAV9-Cre animals. On average, AAV-injected hemispheres had summed striatal areas of editing of 1.40 ± 0.33 mm^2^ compared to NC-Ai14 hemispheres at 1.69 ± 0.72 mm^2^, though one outstanding NC-Ai14 injected hemisphere surpassed this grouping with an area of 7.39 mm^2^. The mean striatal edited area in the NC-Ai14 group without the outlier was 0.977 ± 0.12.

**Figure 2.**
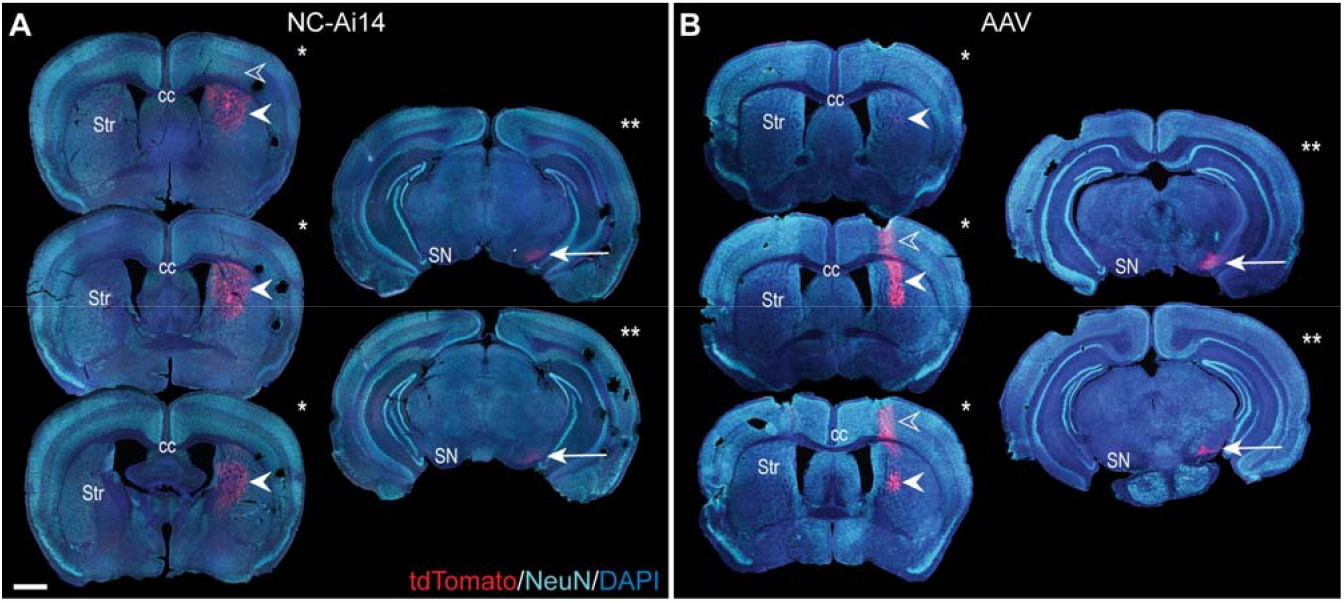
The rostrocaudal distribution of the reporter protein tdTomato in an NC-Ai14-treated hemisphere (A) and an AAV9-Cre-treated hemisphere (B). TdTomato is visible in the cortex along the needle tract in a small number of sections (open arrowheads). TdTomato is abundantly present at the striatal (Str) injection site (filled arrowheads); the three largest striatal areas of editing for each hemisphere are denoted by a single asterisk. Within the substantia nigra (SN), expression of tdTomato is observable (arrow); the two tissue sections with the largest area of tdTomato in the SN are denoted by two asterisks. Scale bar = 1000 μm. AAV, adeno-associated virus; DAPI, 4′,6-diamidino-2-phenylindole; NeuN, neuronal nuclear protein.

TdTomato expression was mainly observed in neuron-like cells characterized by triangular-shaped cell bodies and co-labeled with the neuronal marker NeuN (Figure 3A, C). The intense tdTomato fluorescence was observed in the soma of these cells, with lighter fluorescence signal in the fibers emanating from the soma, corresponding to axons and dendrites. A lower density of edited neurons was visible in hemispheres injected with NCs, a phenomenon that appeared to be coupled with downregulation of NeuN in the areas of injection (Figures 2A and 3A). In addition to the higher density of tdTomato+ neuronal cells in AAV-injected hemispheres, a higher intensity in the red channel was observed broadly throughout the injection site compared to NC-Ai14 hemispheres, possibly due to a higher density of tdTomato+ axons and dendrites (Figure 3C).

**Figure 3.**
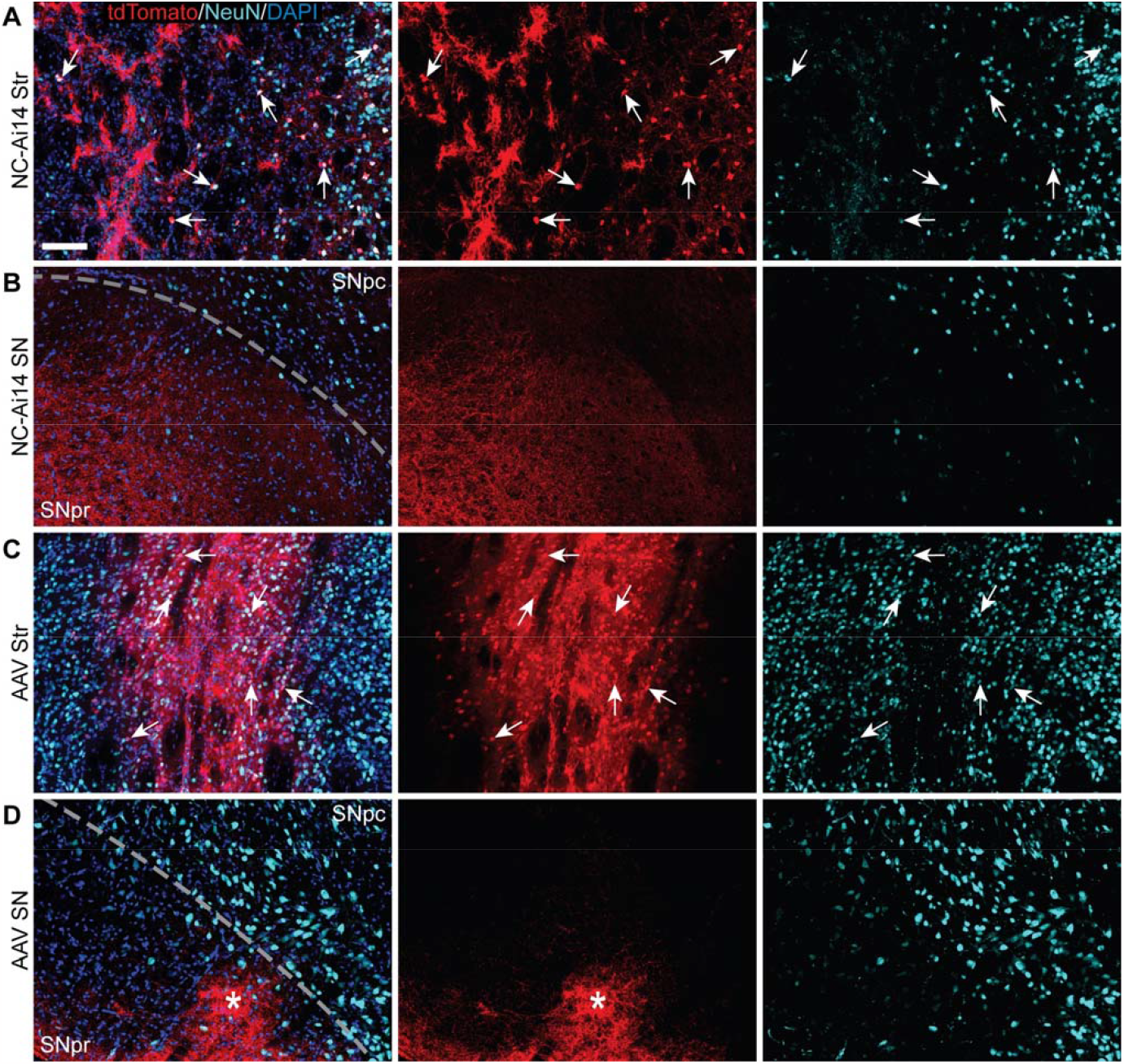
TdTomato expression in striatal and nigral ROIs in a NC-Ai14-injected hemisphere (A, B) and an AAV9-Cre-injected hemisphere (C, D). Within the striatum (Str; A, C), a high degree of colocalization of tdTomato (red) with the neuron marker NeuN (light blue) was observed, indicating successful neuron gene editing (arrows). Within the substantia nigra (SN) ipsilateral to the injected striatum (B, D), tdTomato expressing fibers are clearly visible in the SNpr and often organized into defined bundles of fibers (asterisks). Scale bar = 100 μm. AAV, adeno-associated virus; DAPI, 4′,6-diamidino-2-phenylindole; NeuN, neuronal nuclear protein.

### 3.2 Expression of tdTomato was observed in the SNpr and rarely in the SNpc

TdTomato-ir was visible in both NC-Ai14 and AAV9-Cre treated animals in the SNpr ipsilateral to the injected striatum (Figures 2 and 3). The expression of tdTomato followed along the long, spindly axonal fibers and was not observed in cell bodies in the SNpr (Figure 3B, D). In nearly all sections analyzed, this axonal tdTomato-ir was distributed heterogeneously across the SNpr, appearing in discrete clusters of fibers (Figure 3D). This confined pattern of expression in SNpr fibers was not observed in the NC-injected hemisphere with an outstandingly large area of striatal editing, where tdTomato+ fibers were observed diffusely throughout most of the SNpr in this hemisphere (Figure 3B). The average summed area of the SNpr containing tdTomato+ fibers appeared higher in the AAV9-Cre group (0.228 ± 0.072 mm^2^) than in the NC-Ai14 group (0.168 ± 0.060 mm^2^), although this difference did not reach statistical significance. The mean SNpr tdTomato+ area in the NC-Ai14 group without the animal with notably large striatal edited area was 0.109 ± 0.017 mm^2^. It should also be noted that hemispheres injected with AAV vectors displayed higher intensity of fluorescence signal in the red channel in the SNpr than hemispheres injected with NCs (Figure 3 B, D), similar to observations in the striatum.

In the SNpc, tdTomato expression was rarely observed. In one AAV9-Cre animal, a cluster of three TdTomato+ cells were visible in the SNpc ipsilateral to the injection and co-labeled for the dopaminergic neuron marker tyrosine hydroxylase (TH) (Figure 4; arrowheads).

**Figure 4.**
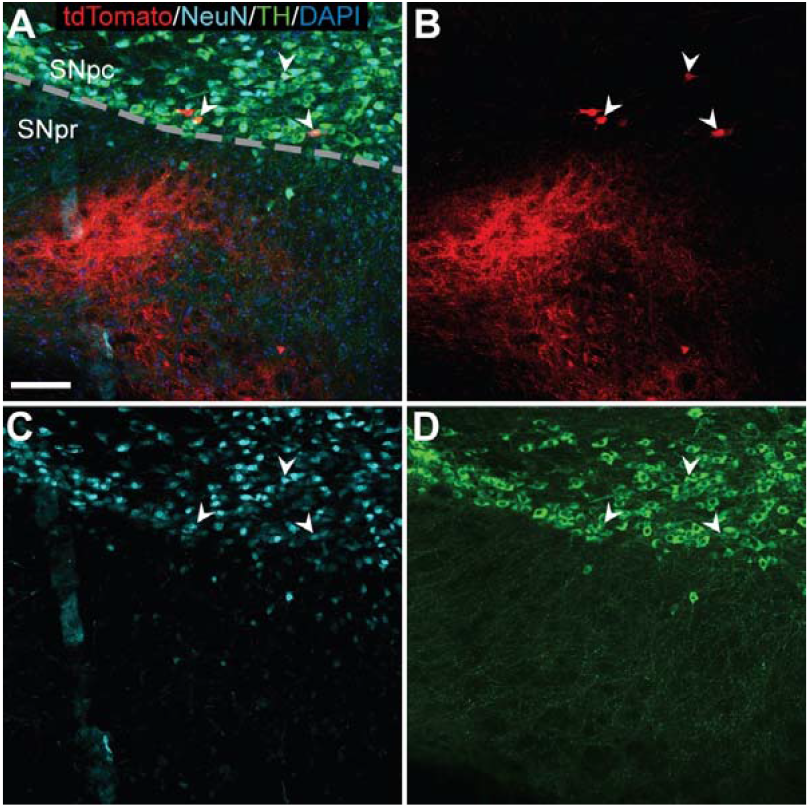
TdTomato expression in TH+ nigral neurons of an AAV9-Cre injected hemisphere. Triple label immunofluorescence (A) identified few tdTomato+ cells (B; red) that colocalized with NeuN (C; light blue) and TH (D) (arrowheads) indicating successful gene editing of dopaminergic neurons in the substantia nigra pars compacta (SNpc). Scale bar = 100 μm. DAPI, 4′,6-diamidino-2-phenylindole; NeuN, neuronal nuclear protein; TH, tyrosine hydroxylase; substantia nigra pars reticulata (SNpr).

### 3.3 There is a relationship between tdTomato+ edited area size in the striatum and the SNpr

To investigate the relationship between the size of the tdTomato+ genome edited area in the striatum and the SNpr, we performed a Pearson correlation test. A statistically significant positive correlation was observed (r = 0.919, p < 0.00001; Figure 5A), which remained significant when the NC-Ai14 animal with notably large striatal edited area was excluded (r = 0.810, p < 0.00001; Figure 5B). This analysis included hemispheres that received striatal NC-Scrambled injections, which exhibited no editing in the striatum and minimal background signal in the SNpr.

**Figure 5.**
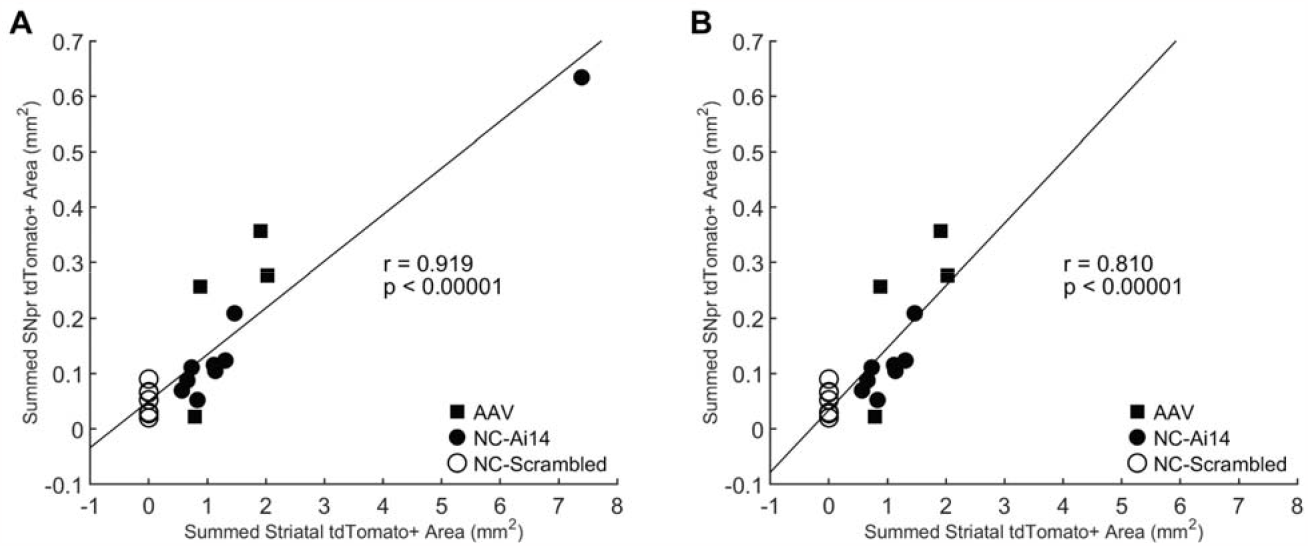
Correlation between the size of the tdTomato+ genome edited area in the striatum and SNpr. A statistically significant correlation was observed for Ai14 mice injected with NC-Ai14 (n=9), NC-Scrambled (n=9), or AAV9-Cre (n=4) (A), which remained significant when the NC-Ai14 animal with notably large striatal edited area was excluded (B). AAV, adeno-associated virus; NC, nanocapsule.

## 4 Discussion

Overall, these results demonstrate that *in vivo* neuronal genome editing in the murine striatum produces changes in reporter gene expression in anatomically connected regions of the striato-nigral neural network. Both NC-Ai14 and AAV9-Cre injections led to successful gene editing in Ai14 mouse striatal neurons, and the size of the area of striatal editing correlated with the amount of tdTomato expression in the ipsilateral SNpr. Editing of neuronal DNA in the striatum was required for visualization of tdTomato in the SNpr, as no animals in the NC-Scrambled group exhibited tdTomato expression in the SN. Approximately 95% of neurons in the striatum are medium spiny neurons (Arlotta et al., 2005), which send inhibitory projections to the surrounding nuclei of the basal ganglia, including the SNpr (Stephenson-Jones et al., 2012; Partanen and Achim, 2022). Our findings suggest that the tdTomato protein produced in the soma of genome edited striatal medium spiny neurons undergoes axonal anterograde transport to fill the cell cytoplasm and appear visible in fibers in the SNpr.

Expression of tdTomato in the SNpr was typically present in discrete clusters of fibers and limited to animals with total areas of striatal editing above approximately 1 mm^2^. The nonuniform distribution of tdTomato in the SNpr is consistent with previous descriptions of striatal inputs to the SNpr maintaining features of the striatal topographical organization (Partanen and Achim, 2022). Tracing studies in rodents have shown that axons from GABAergic medium spiny neurons in the striatum are directed to defined longitudinal columns across the SNpr (Deniau et al., 1996; Foster et al., 2021). Related to this, the NC treated animal with more diffuse presence of tdTomato in the SNpr also showed the largest area with genome edited neurons in the striatum. It is currently unclear why animals with small striatal edited areas below 1 mm^2^ did not have tdTomato present in SNpr fibers. It is possible that, if tdTomato was present in the SNpr in these animals, it was in too few axons and/or too dim to be detectable above background signal in the red channel.

In a single AAV9-Cre treated animal, a small cluster of tdTomato-filled neuronal cell bodies was present in the SNpc. Given the well-characterized anatomical connection between the dopaminergic SNpc neurons and the striatum, these tdTomato+ nigral neurons likely represent uptake of AAV9-Cre by SNpc terminals in the striatum followed by retrograde transport of the viral vector to the nigral soma. Retrograde transduction of nigral neurons following striatal AAV9 delivery has been previously described in in rats (Löw et al., 2013), although this publication indicates that another AAV serotype, AAV6, produced significantly greater efficiency of retrograde transduction. Additionally, a study in nonhuman primates demonstrated both anterograde and retrograde transportation of AAV9 carrying green fluorescent protein injected into the striatum (Green et al., 2016). It has been reported that AAV9 can enter neuronal fibers and undergo retrograde transport within late endosome compartments using dynein/dynactin (Castle et al., 2014a; Castle et al., 2014b). NCs have been shown to colocalize with endosomes/lysosomes immediately after transfection and their RNP payload to colocalize within nuclei five hours after transfection (Metzger et al., 2023), however, mechanisms underlying potential axonal transport have not yet been evaluated. Although no tdTomato+ SNpc neurons were observed in NC treated animals in this study, future experiments focused on mechanisms of NC neuronal anterograde and retrograde transport are warranted.

Although both NC-Ai14 and AAV9-Cre both produced neuronal genome editing, it was observed throughout confocal microscope image acquisition that red channel signal intensity was noticeably higher in AAV injected animals (see methods). Increased tdTomato intensity in AAV animals is potentially related to the mechanisms of editing the Ai14 reporter allele. AAV9-induced Cre-lox recombination removes the entirety of the stop cassette upstream of the tdTomato reporter gene sequence. In contrast, the guide RNA in the NC-Ai14 CRISPR RNP targets repeated sequences between the SV40 polyA transcriptional terminator blocks within the cassette, and excision of at least two of the three SV40 polyA blocks is required to produce tdTomato (Staahl et al., 2017; Metzger et al., 2023). Because of this, expression of the tdTomato protein is known to underreport total NC gene editing, and cells with incomplete removal of the stop cassette may have shown reduced expression of tdTomato relative to cells transduced with AAV9-Cre. Furthermore, while previous work has shown that intrastriatal NC injection led to primarily neuronal editing (Metzger et al., 2023), AAV9 shows broader tropism across multiple cell types in the nervous system, particularly astrocytes (Foust et al., 2009; Gray et al., 2011; Hammond et al., 2017). The presence of tdTomato in genome edited astrocytes in the striatum may have further contributed to red channel signal intensity in AAV injected animals.

Successful application of somatic genome editing for the treatment of central nervous system disorders will necessarily be impacted by the anatomy of the highly distributed neural networks of the brain. Disease pathophysiology can further affect the distribution and efficacy of delivered therapies. For example, despite abundant preclinical evidence for neuroprotective effects of glial cell line-derived neurotrophic factor (GDNF) delivered to the striatum or the SN in animal models of PD, clinical trials in patients with PD have largely failed. Although there is debate as to why clinical translation has been unsuccessful, the degeneration of the SNpc dopaminergic neurons and their axons in patients could prevent response to growth factor therapy (Manfredsson et al., 2020; Kambey et al., 2021). An important next step in testing the potential application of NC or AAV delivery of gene editors to address human neurological disease will be evaluation of distribution of editors across neural networks in large animal models, such as nonhuman primates, and in models of diseases.

## 6 Author Contributions

SSN conceptualized, drafted, and revised the article and was involved in methodology development, investigation, visualization, and formal analysis. JMM conceptualized, drafted, and revised the article and was involved in methodology development and investigation. VB, YW, and JF were involved in formal analysis and investigation. JEL, KS, and SG conceptualized and revised the article as well as provided resources, supervision, project management, and funding acquisition. MEE conceptualized, drafted, and revised the article and was involved in methodology development, validation, data curation, supervision, project management, and funding acquisition.

## 8 Acknowledgments

We are grateful for the technical support provided by Kat Jones, Pankaj Dubey, Matthew T. Flowers, and Kirsten Gimse.

## 9 Data Availability Statement

The data that support the findings of this study are available from the corresponding author upon reasonable request.

